# Lysosomes Signal through Epigenome to Regulate Longevity across Generations

**DOI:** 10.1101/2025.05.21.652954

**Authors:** Qinghao Zhang, Weiwei Dang, Meng C. Wang

## Abstract

Epigenome is sensitive to metabolic inputs and crucial for aging. Lysosomes emerge as a signaling hub to sense metabolic cues and regulate longevity. We unveil that lysosomal metabolic pathways signal through the epigenome to regulate transgenerational longevity in *Caenorhabditis elegans*. We discovered that the induction of lysosomal lipid signaling and lysosomal AMP-activated protein kinase (AMPK), or the reduction of lysosomal mechanistic-target-of-rapamycin (mTOR) signaling, increases the expression of histone H3.3 variant and elevates H3K79 methylation, leading to lifespan extension across multiple generations. This transgenerational pro-longevity effect requires intestine-to-germline transportation of H3.3 and a germline-specific H3K79 methyltransferase, and can be recapitulated by overexpressing H3.3 or the H3K79 methyltransferase. This work uncovers a lysosome-epigenome signaling axis linking soma and germline to mediate the transgenerational inheritance of longevity.

## Main Text

Lysosomes are pivotal in sensing nutrient availability in the environment. During the period of starvation, the lysosomal LIPL-4 lipase is transcriptionally upregulated, lysosomal recruitment of AMPK is enhanced, and lysosomal mTOR activation is suppressed (*1-3*). LIPL-4, AMPK and mTOR signaling pathways as well as starvation, have all been linked to longevity regulation (*4*). Starvation has also been reported to induce epigenetic inheritance across generations in diverse organisms (*5*, *6*). However, it remains unclear whether and how those lysosomal signaling pathways formulate epigenetic memories and contribute to transgenerational longevity. In this study, we discovered a soma-to-germline epigenetic axis that is activated in response to lysosome-related signaling pathways and promotes longevity across generations.

### Lysosomal lipolysis elevates H3K79 methylation and promotes transgenerational longevity

LIPL-4 is a lysosomal acid lipase, the overexpression of which in the intestine, the major metabolic tissue of worms, induces lysosomal lipolysis and promotes longevity (*7*, *8*). When back crossing the long-lived *lipl-4* transgenic (*lipl-4 Tg*) worms with wild-type (WT) worms, we noticed that the WT descendants from the cross show extended lifespan, suggesting that the LIPL-4-induced lysosomal lipolysis may promote longevity across generations. To confirm this transgenerational pro-longevity effect, we crossed *lipl-4 Tg* hermaphrodites with WT males and followed F3 WT descendants, designated as second transgenerational WT (T2 WT), for lifespan analyses (Fig. 1A). In parallel, WT hermaphrodites were crossed with WT males and F3 progeny were used as WT controls (Fig. 1A). We found that T2 WT descendants from the *lipl-4 Tg* cross live longer than WT controls (Fig. 1A and table S1).

**Fig. 1.**
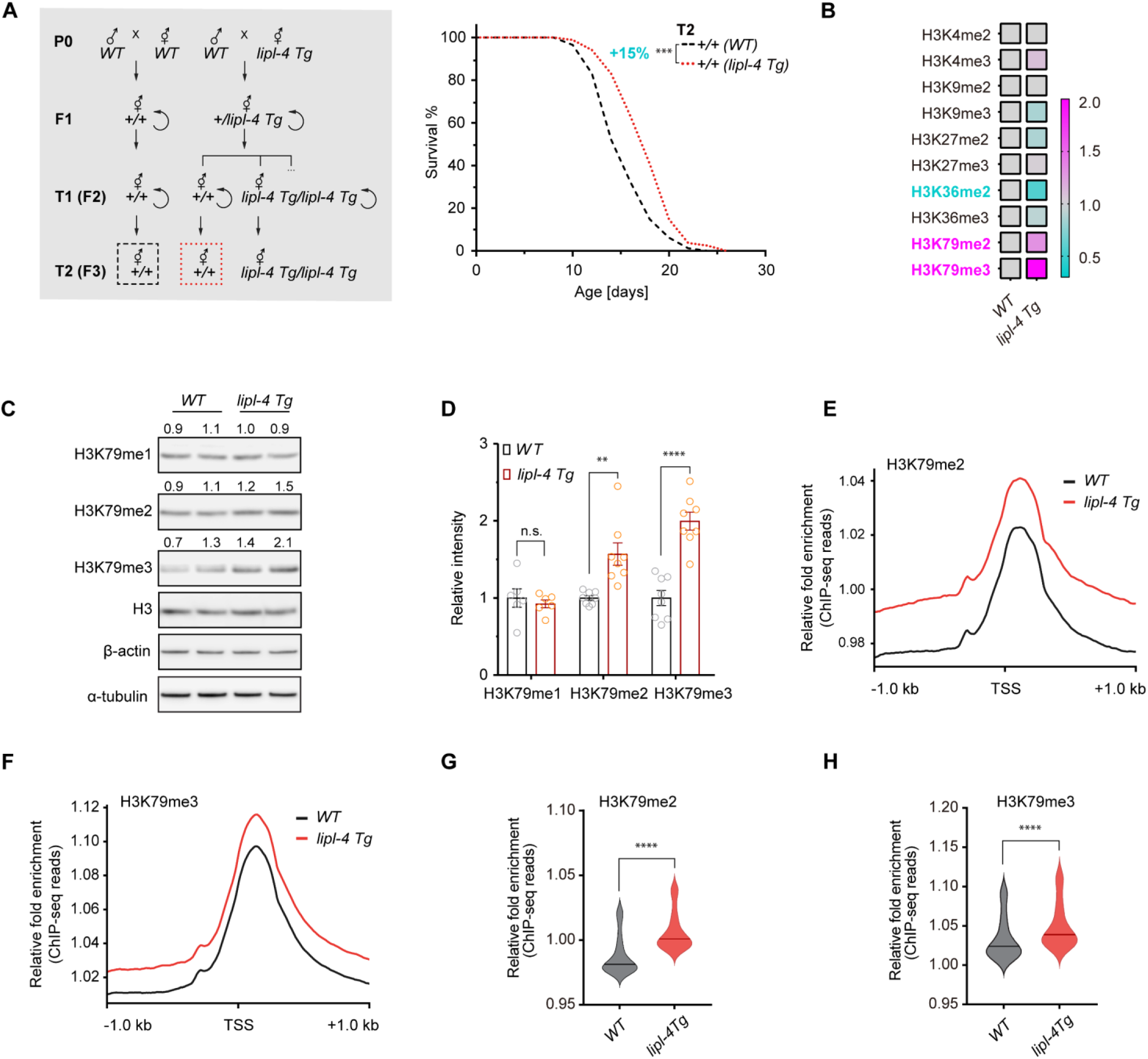
Lysosomal lipolysis induces transgenerational inheritance of longevity and H3K79 methylation. (**A**) T2 WT descendants (*+/+ (lipl-4 Tg)*), which are second transgenerational WT from P0 *lipl-4 Tg* worms, exhibit extended lifespan compared to WT controls (*+/+ (WT)*) from P0 WT worms. With n = 90/replicate, 5 biological replicates; log-rank test followed by Fisher’s method: \*\*\**p* < 0.001. Percentage of lifespan extension (lower vs. upper) is labeled. Summary of lifespan replicates shown in table S1. Scheme describing genetic crosses and transgenerational groups (framed with dotted boxes) used in lifespan analyses. (**B**) The heatmap reveals the ratio of histone H3 PTM levels in *lipl-4 Tg* vs. WT worms from Western blot screens. (**C**) Western blot images show levels of H3K79 mono-(me1), di-(me2) and tri-methylation (me3), histone H3, β-actin and ⍺-tubulin in WT and *lipl-4 Tg* worms. (**D**) The bar chart summarizes all biologically independent replicates from Western blots. Error bars represent mean ±standard error of the mean (s.e.m.), n.s. *p* > 0.05, \*\**p* < 0.01, \*\*\*\**p* < 0.0001 (unpaired t-test, Welch’s correction). (**E-H**) Enhanced deposition (±1 kb centered over TSS) of both H3K79me2 and H3K79me3 was observed in *lipl-4 Tg* vs. WT worms, which is quantified in violin plots (central lines denote median values). \*\*\*\**p* < 0.0001 (Wilcoxon test).

Epigenetic changes play crucial roles in the regulation of transgenerational inheritance. In particular, H3K4me3 and H3K9me2 have been associated with transgenerational longevity in *C. elegans* (*9*, *10*). We found that neither H3K4me3 nor H3K9me2 exhibits a profound alteration in the *lipl-4 Tg* worms (Fig. 1B and fig. S1A). We then compared the levels of other histone H3 methylations, including H3K4me2, H3K9me3, H3K27me2, H3K27me3, H3K36me2, H3K36me3, H3K79me2 and H3K79me3, between the *lipl-4 Tg* and WT worms at day-1 adulthood (Fig. 1B and fig. S1A). We found that the levels of H3K79me2 and H3K79me3 are both increased in the *lipl-4 Tg* worms (Fig. 1B-D, and fig. S1).

To validate the upregulation of H3K79me2 and H3K79me3, we further conducted the chromatin immunoprecipitation followed by next-generation sequencing (ChIP-seq) using anti-H3K79me2 and anti-H3K79me3 antibodies and day-1 adult WT and *lipl-4 Tg* worms. Through metaprofile analysis, both H3K79me2 and H3K79me3 marks were mounted at peaks prior to the transcription start sites (TSS) and showed enrichments within the proximal transcribed region of gene bodies (Fig. 1E, 1F), which is consistent with previous reports in mammals (*11*). In comparison with the WT genome, the *lipl-4 Tg* genome exhibited increased occupancy of both H3K79me2 and H3K79me3, spanning 1 kb downstream and upstream of TSS (Fig. 1E-H). These results support the Western blot results that H3K79me2 and H3K79me3 levels were increased in the *lipl-4 Tg* worms.

### Lysosomal lipolysis transcriptionally upregulates H3.3 variant HIS-71

It is known that H3K79 methylation commonly serves as a mark for transcriptional activation (*12*, *13*). To further decipher the molecular target that initiates the transgenerational pro-longevity effect, we integrated the H3K79me2/me3 ChIP-seq data with the RNA-seq data from the *lipl-4 Tg* worms, with a focus on those transcriptionally upregulated genes. From the ChIP-seq data, we found 1469 sites showing statistically differential occupancy of H3K79me2 between the *lipl-4 Tg* and WT worms (Fig. 2A), and 2335 sites showing statistically differential occupancy of H3K79me3 (Fig. 2B) (*p* < 0.01, FDR < 0.05). After mapping all these differential sites into genes, 500 genes showed enrichments of both H3K79me2 and H3K79me3 in the *lipl-4 Tg* worms (Fig. 2C), with 42 of them transcriptionally upregulated in the RNA-seq analysis (fold change > 1.5, *p* < 0.01, FDR < 0.05) (Fig. 2D).

**Fig. 2.**
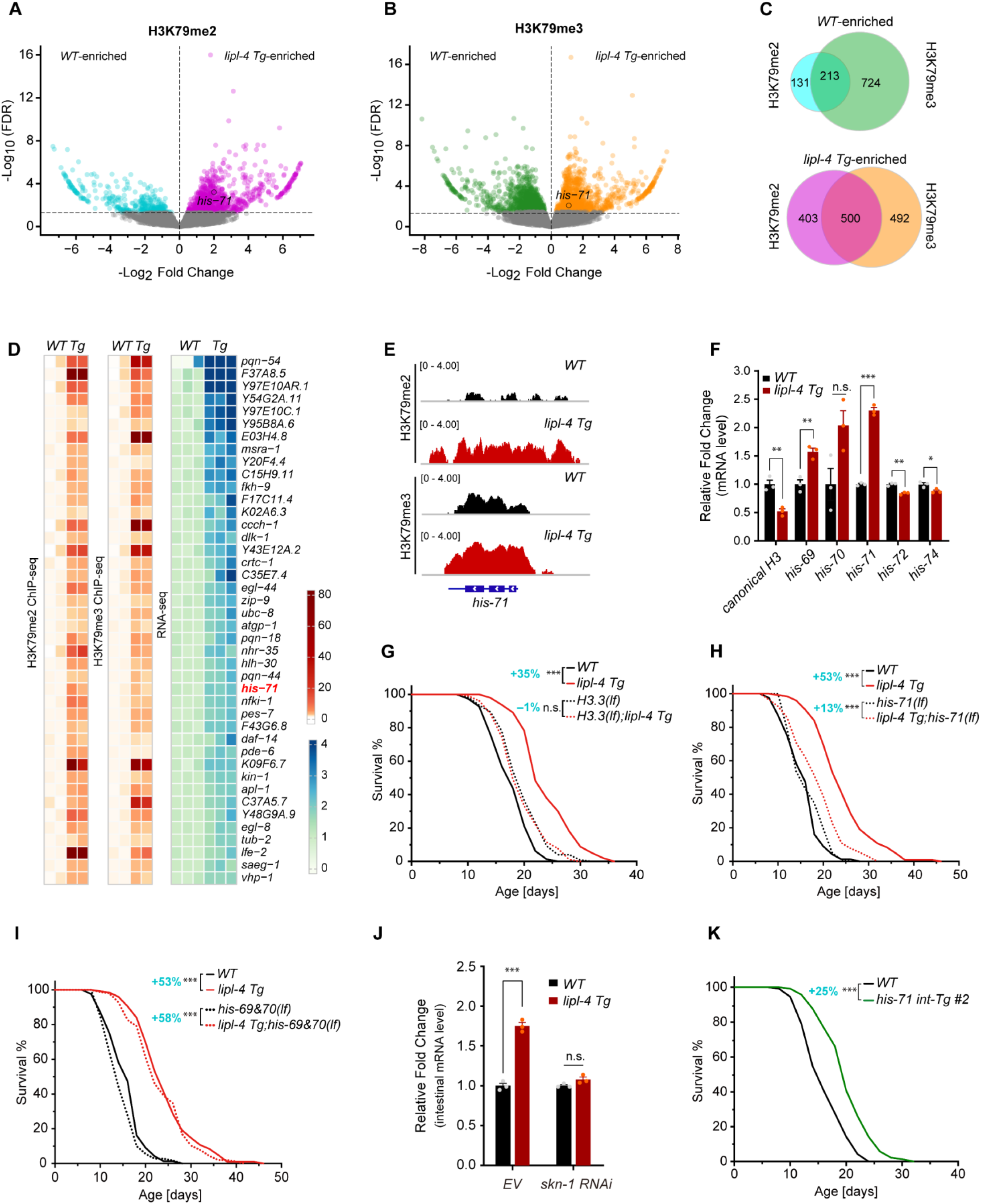
Lysosomal lipolysis transcriptionally upregulates H3.3 variant HIS-71 to promote longevity. (**A, B**) Volcano plots reveal gene loci with differential H3K79me2 or H3K79me3 deposition between WT and *lipl-4 Tg* worms. Cyan/green, higher occupancy in WT; magenta/orange, higher in *lipl-4 Tg*; gray, no significant difference; encircled, *his-71*. (**C**) Venn diagrams show the numbers of genes enriched with H3K79me2 and/or me3 marks specifically in WT or *lipl-4 Tg* worms. (**D**) Heatmaps reveal gene candidates that are enriched with both H3K79me2 and H3K79me3 marks and are transcriptionally upregulated in *lipl-4 Tg* vs. WT worms. (**E**) Both H3K79me2 and H3K79me3 marks are elevated on the promoter region and gene body of *his-71* in *lipl-4 Tg* vs. WT worms. (**F**) Relative expression changes of canonical H3 and H3.3 variants in *lipl-4 Tg* vs. WT worms. (**G**-**I**) The null mutant of all H3.3 variants or the loss-of-function (lf) mutant of *his-71*, but not the *his-60&70(lf)* mutant, suppresses the lifespan extension caused by *lipl-4 Tg*. (**J**) Intestinal *his-71* expression is upregulated in *lipl-4 Tg* vs. WT worms, which is suppressed by *skn-1* RNAi inactivation. (**K**) Intestine-specific overexpression of *his-71* (*his-71 int-Tg*) extends lifespan. In (G-I and K), n = ∼90/replicate, 3 biological replicates; log-rank test followed by Fisher’s method: n.s., *p* > 0.05, \*\*\**p* < 0.001. Percentage of lifespan extension (lower vs. upper) is labeled. Summary of lifespan replicates shown in table S2. In (F and J), error bars represent mean ±s.e.m., n.s., *p* > 0.05, \**p* < 0.05, \*\**p* < 0.01, \*\*\**p* < 0.001 (unpaired t-test, Welch’s correction).

Interestingly, one of these targets is *his-71* (Fig. 2D, 2E), which encodes a histone variant H3.3. As a key epigenetic regulator of gene expression networks, histone H3.3 is incorporated into chromatin independently of replication in differentiated cells (*14-16*), and displays a greater propensity for K79 methylation than canonical H3 (*17*, *18*). The *C. elegans* genome contains five H3.3 encoding genes, *his-69*, *his-70*, *his-71*, *his-72,* and *his-74* (*19*, *20*), and 15 genes encoding canonical H3 (*21*). We found that the occupancy of H3K79me2/me3 on the other H3.3 variant genes or any canonical H3 genes are not elevated in *lipl-4 Tg* vs. WT worms (fig. S2A). When examining their transcriptional levels by qRT-PCR, *his-71* and *his-69* were transcriptionally upregulated, while canonical H3 was down-regulated in the *lipl-4 Tg* worms (Fig. 2F). The induction of *his-70* was ∼2-fold, although it did not reach statistical significance (*p* > 0.05, Fig. 2F). We further investigated the role of H3.3 variants in the regulation of longevity. We found that the null mutation of H3.3 reduces the lifespan extension (from +35% to −1%) caused by *lipl-4 Tg* (Fig. 2G and table S2), so does the *his-71* deletion (*his-71(lf)*) (from +53% to +13%) (Fig. 2H and table S2). On the other hand, deletion of *his-69* and *his-70* did not suppress the lifespan extension in the *lipl-4 Tg* worms (Fig. 2I and table S2).

LIPL-4 acts in the intestine to promote longevity (*7*, *8*). We further confirmed that *his-71* is transcriptionally upregulated in the intestine of the *lipl-4 Tg* worms (Fig. 2J, fig. S2B). In our previous studies, we have identified several transcription factors that mediate the pro-longevity effect in the *lipl-4 Tg* worms, including NHR-49, NHR-80, JUN-1, and SKN-1 (*8*, *22*). When performing a bioinformatic analysis of the promoter regions from the 42 candidate genes, we found that 36 of them have promoter regions longer than 1 kb, and the SKN-1 binding motif is enriched in the promoter region of 18 candidate genes, including *his-71* (fig. S2C). Consistently, *skn-1* RNAi knockdown suppresses the transcriptional upregulation of *his-71* in the intestine of the *lipl-4 Tg* worms (Fig. 2J). Furthermore, we found that intestine-specific overexpression of *his-71* (*his-71 int-Tg*) sufficiently prolongs lifespan (Fig. 2K, fig. S2D, table S2). Together, these results suggest that LIPL-4-induced lysosomal lipolysis upregulates the expression of the H3.3 variant *his-71* in the intestine to promote longevity.

### HIS-71 transports to germline and mediates transgenerational longevity

Interestingly, when assessing intestinal *his-71* transcriptional upregulation across generations in WT descendants from the *lipl-4 Tg* worms, we observed a progressive decline, with the upregulation completely lost by T4 generation (Fig. 3A). This concords with the progressive loss of the longevity promoting effect from T1 to T4 WT generations (fig. S3A-C, Fig. 1A and table S1), suggesting a potential causal relationship between the transgenerational longevity and *his-71* upregulation. We thus examined whether HIS-71 mediates transgenerational longevity and found its deletion fully suppresses the lifespan extension in the T2 WT descendants from the *lipl-4 Tg* cross (Fig. 3B, fig. S3D, and table S1). Furthermore, we generated a CRISPR knock-in line in which the endogenous HIS-71 is tagged with a degron, and then crossed this line with the *lipl-4 Tg* worms, as well as with the transgenic strain where the auxin-inducible F-box protein TIR1 is selectively expressed in the intestine. Using the crossed strain, we could induce TIR1-mediated degradation of the degron-tagged HIS-71 protein selectively in the intestine by the auxin treatment (fig. S3E). We administered auxin to T1 WT worms from the *lipl-4 Tg* cross and found that the intestine-specific HIS-71 degradation suppresses the lifespan extension in the T2 WT descendants (Fig. 3C, fig. S3F, and table S1). We also confirmed that selectively overexpressing *his-71* in the intestine promotes transgenerational longevity (Fig. 3D and table S1). These results suggest that intestinal HIS-71 contributes to the transgenerational pro-longevity effect conferred by lysosomal lipolysis.

**Fig. 3.**
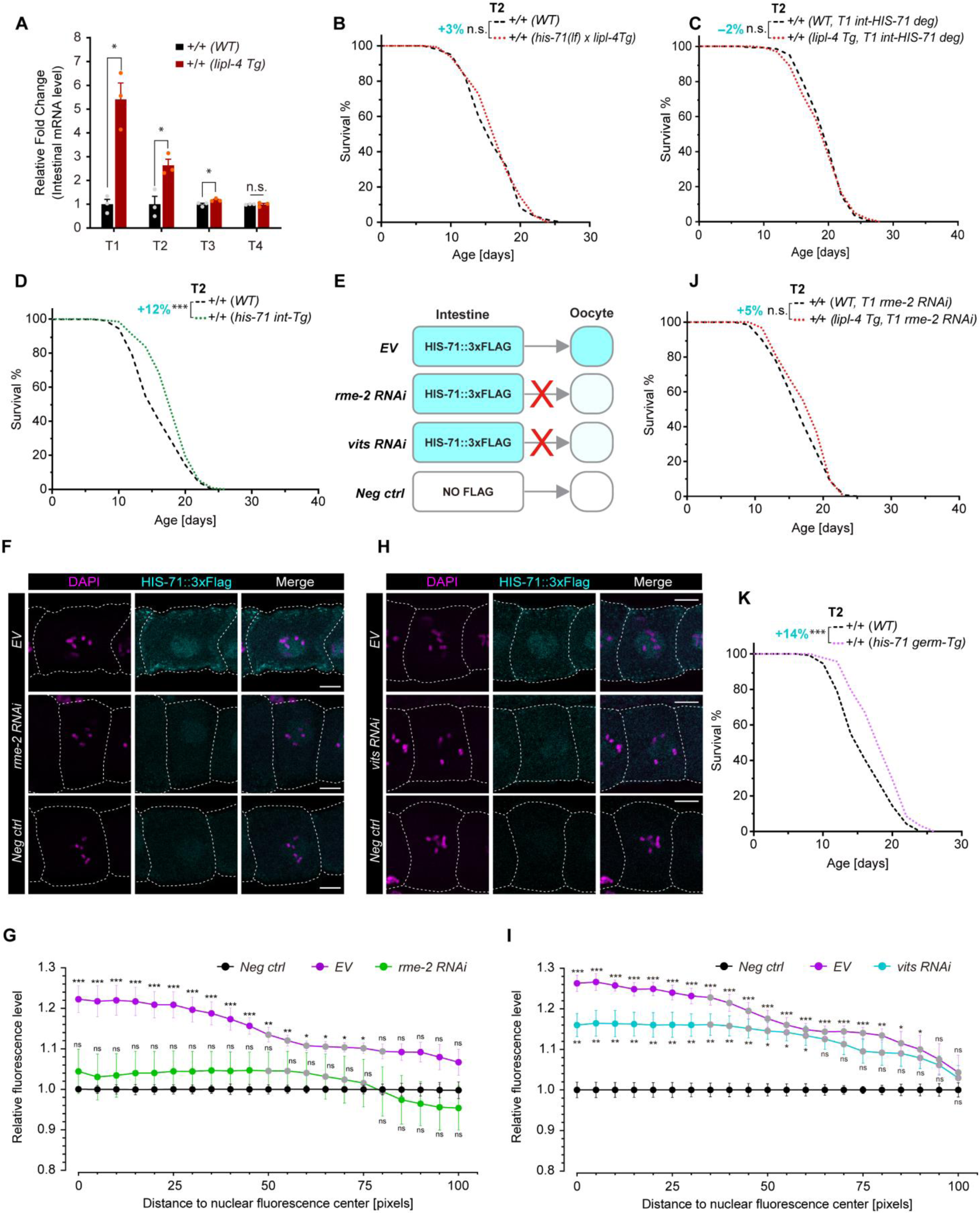
Intestinal HIS-71 and its transportation to germline contributes to transgenerational longevity. (**A**) Intestinal upregulation of *his-71* transcription in WT descendants of *lipl-4 Tg* vs WT worms declines across generations. Error bars represent mean ±s.e.m., n.s., *p* > 0.05, \**p* < 0.05 (unpaired t-test, Welch’s correction). (**B**)The *his-71(lf)* mutation abrogates the transgenerational longevity in T2 WT from *lipl-4 Tg*. (**C**) Upon intestine-specific degradation of HIS-71 in the T1 generation (T1 int-HIS-71 deg), the transgenerational longevity caused by *lipl-4 Tg* fails to be transmitted into the T2 generation. (**D**) Intestinal overexpression of *his-71* leads to transgenerational longevity in T2 WT progeny. (**E**-**I**) Intestine-derived HIS-71 is transported into oocytes within the germline, a process that is suppressed by RNAi inactivation of *rme-2* (germline endocytic receptor) (F, G) and *vitellogenin* genes (*vits*) (H, I). Images represent oocytes at the proximal −2 position from germline immunostaining using anti-FLAG antibody. Worms with intestine-specific overexpression of *his-71::3×flag* were subjected to the empty vector control (EV) and *rme-2* or *vits* RNAi, and worms with intestine-specific overexpression of *his-71* without *3×flag* served as a negative control (Neg ctrl). Scale bar = 10 μm (97.5 pixels). (G, I) Quantification of anti-FLAG fluorescent signals in −2 oocytes, n = 7-15 per replicate, 4 biological replicates. Error bars represent mean ± s.e.m. Two-way ANOVA with Bonferroni’s test: n.s., *p* > 0.05, \**p* < 0.05, \*\**p* < 0.01, \*\*\**p* < 0.001 (EV/ *rme-2*/ *vits* RNAi vs. Neg ctrl); gray dots, *p* > 0.05, otherwise, *p* < 0.05 (*rme-2*/ *vits* RNAi vs. EV). (**J**) RNAi inactivation of *rme-2* in the T1 generation (T1 *rme-2* RNAi) abolishes the transmission of LIPL-4-induced transgenerational longevity to T2. (**K**) Germline overexpression of *his-71* (*his-71 germ-Tg*) promotes transgenerational longevity in T2 WT progeny. In (B-D, J, K), n = 90/replicate, 3 biological replicates; log-rank test followed by Fisher’s method: n.s., *p* > 0.05, \*\*\**p* < 0.001. Percentage of lifespan extension (lower vs. upper) is labeled. Summary of lifespan replicates shown in table S1.

Transgenerational inheritance requires the transmission of epigenetic information through the germline across generations. Given the HIS-71’s impact on transgenerational longevity, we questioned whether this H3.3 variant could be transferred from the intestine to the germline. To investigate, we generated a transgenic strain that exclusively expresses HIS-71 tagged with 3×FLAG in the intestine, without ectopic expression in the germline (table S3). We dissected the germline from this strain to conduct immunostaining using anti-FLAG antibodies (Fig. 3E). The HIS-71::3×FLAG, originating from the intestine, was detected in the oocytes within the germline (Fig. 3F-I). Vitellogenins are large lipid transfer proteins synthesized in the intestine and transported into oocytes via RME-2, a lipoprotein-like endocytic receptor, and serve as a nutrient source for embryos (*23*, *24*). We found that the observed HIS-71::3×FLAG signal in the germline was abolished by RNAi knockdown of *rme-2* and reduced by RNAi knockdown of vitellogenin genes (Fig. 3E-I). We also confirmed that the *rme-2* knockdown did not affect the transcriptional induction of *his-71* in the intestine of the *lipl-4 Tg* strain (fig. S3G). These results suggest the transportation of HIS-71 from the intestine to the germline through the vitellogenin secretion and uptake pathway.

Importantly, when knocking down *rme-2* to block the uptake of HIS-71 into the germline of T1 WT derived from the *lipl-4 Tg* worms, we found it is sufficient to abrogate the lifespan extension in the T2 WT descendants (Fig. 3J and table S1), suggesting that the transportation of HIS-71 from the intestine to the germline is required for the transgenerational pro-longevity effect. Furthermore, we induced selective degradation of HIS-71 in the germline using the auxin treatment (fig. S3H), and this degradation in the T1 WT descendants from the *lipl-4 Tg* cross fully abolished the lifespan extension in the T2 WT descendants (fig. S3I and table S1). Conversely, germline-specific overexpression of *his-71* (*his-71 germ-Tg*) promotes longevity across generations (Fig. 3K, fig. S3J, S3K, and tables S1, S2). Thus, the local upregulation of *his-71* is sufficient to promote transgenerational longevity. Interestingly, when analyzing *his-71* transcription in the germline, we found that it is upregulated in the *lipl-4 Tg* strain (fig. S3L). We also found that the intestine-specific overexpression of FLAG-tagged *his-71* leads to the upregulation of endogenous *his-71* gene in the germline, which is abolished by hampering HIS-71 intestine-to-germline transportation via RNAi knockdown of *rme-2* or vitellogenin genes (fig. S3M). Together, these results suggest that in the *lipl-4 Tg* worms, transcriptional induction of *his-71* in the intestine would increase the level of the HIS-71/H3.3 in the germline, leading to transgenerational longevity.

### Germline H3K79 methyltransferase promotes longevity

Meanwhile, we also examined whether the increased H3K79 methylation contributes to the transgenerational pro-longevity effect. It is known that H3K79 methylation is added by evolutionarily conserved H3K79 specific N-methyltransferase, Dot1/DOT1L (*25*). *C. elegans* carries six putative H3K79 methyltransferases: DOT-1.1 to −1.5, and D1053.2 (*26*). We knocked down the encoding gene of each DOT1 methyltransferase using RNAi and examined whether any RNAi knockdown reduces the level of lifespan extension in the *lipl-4 Tg* compared to WT worms. We found that upon *dot-1.3* and *dot-1.1* RNAi knockdown, the lifespan extension level in *lipl-4 Tg* vs. WT worms is reduced by ∼30% and ∼20%, respectively, while RNAi knockdown of *dot-1.2, dot-1.4, dot-1.5* or *D1053.2* does not show such reduction (Fig. 4A, fig. S4A-E and table S4). The lifespan extension was also reduced by ∼20% with the *dot-1.3(lf)* mutant, but not with the *dot-1.1(lf)* mutant (fig. S4F, S4G and table S2). Furthermore, the *dot-1.3* knockdown suppressed the induction of H3K79me2 and H3K79me3 in the *lipl-4 Tg* worms (Fig. 4B, fig. S4H and S4I).

**Fig. 4.**
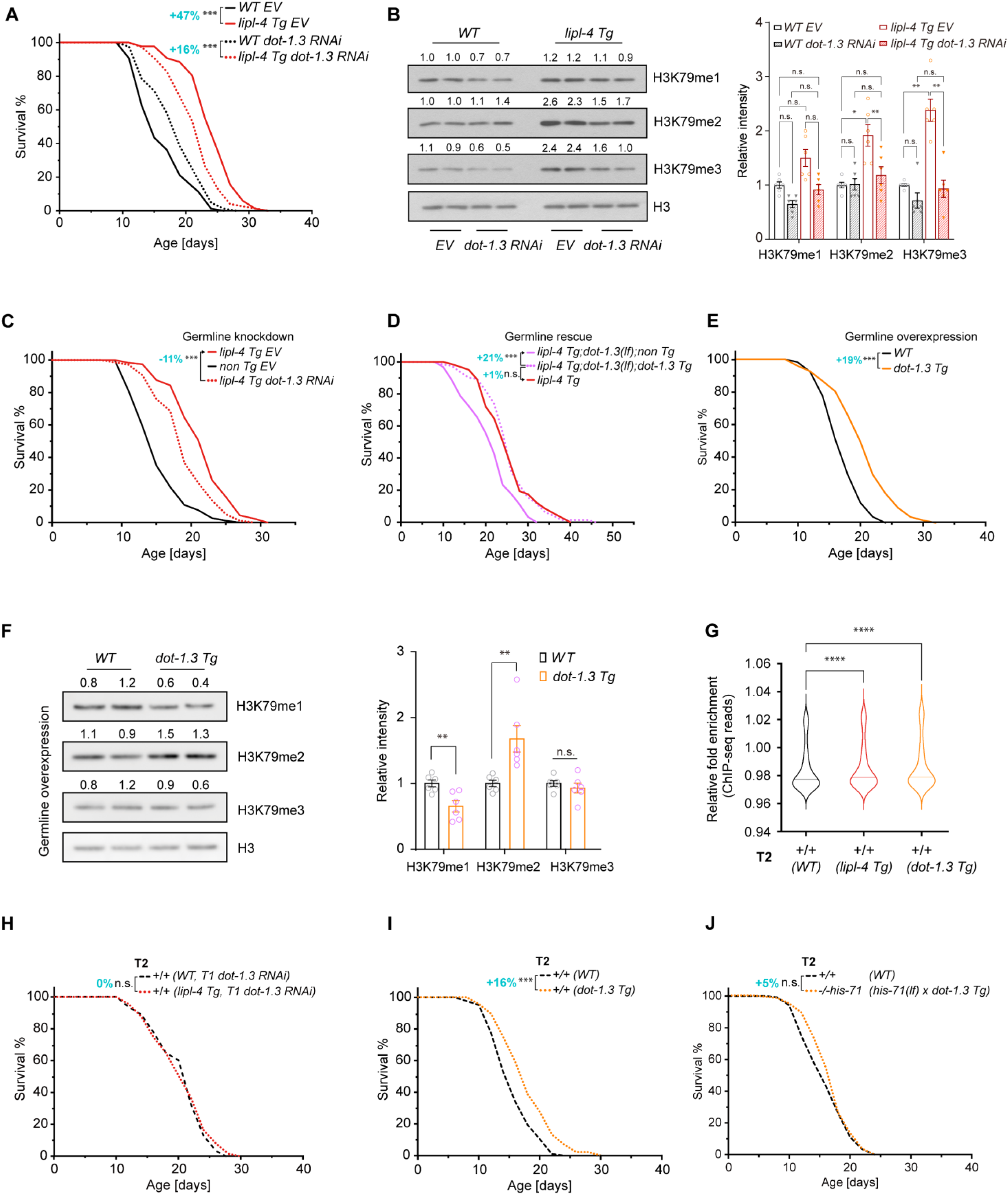
Germline H3K79 methyltransferase DOT-1.3 modulates longevity. (**A**) RNAi inactivation of *dot-1.3* reduces the level of lifespan extension caused by *lipl-4 Tg* from 47% to 16%. (**B**) RNAi inactivation of *dot-1.3* suppresses the induction of H3K79me2/3 in the *lipl-4 Tg* worms. Error bars represent mean ±s.e.m., 6 biological replicates, n.s. *p* > 0.05, \**p* < 0.05, \*\**p* < 0.01 (one-way ANOVA, Tukey test). (**C**) Germline-specific knockdown of *dot-1.3* reduces the lifespan extension caused by *lipl-4 Tg*. (**D**) Rescuing *dot-1.3* expression in the germline restores the lifespan extension in *lipl-4 Tg;dot-1.3(lf)* worms. (**E**) Germline overexpression of *dot-1.3* results in lifespan extension. (**F**) Western blots showing that germline-specific overexpression of *dot-1.3* elevates the H3K79me2 level. Error bars represent mean ± s.e.m., 6 biological replicates, n.s. *p* > 0.05, \*\**p* < 0.01 (unpaired t-test, Welch’s correction). (**G**) T2 WT descendants from *lipl-4 Tg* or *dot-1.3 Tg* exhibit elevated H3K79me2 deposition (±1 kb, TSS) compared to WT descendants from WT. \*\*\*\**p* < 0.0001 (Wilcoxon test). Central lines denote median values of the data. (**H**) RNAi inactivation of *dot-1.3* in the T1 generation (T1 *dot-1.3* RNAi) abrogates the transgenerational longevity in T2 WT progeny from *lipl-4 Tg*. (**I**) T2 WT descendants from *dot-1.3 Tg* show extended lifespan. (**J**) Transgenerational longevity induced by *dot-1.3 Tg* is abolished by *his-71(lf)* mutant. In (A, C-E and H-J), n = ∼90/replicate, 3 biological replicates; log-rank test followed by Fisher’s method: n.s., *p* > 0.05, \*\*\**p* < 0.001. Percentage of lifespan extension (indicated by arrows, or lower vs. upper) is shown. Summary of lifespan replicates shown in tables S1, S2 and S4.

To figure out the tissue-specificity of DOT-1.3, we generated a CRISPR knock-in line in which endogenous DOT-1.3 is tagged with mNeonGreen. We found that DOT-1.3 expression is only detectable in the germline, but not in the intestine or other somatic tissues (fig. S4J). We then knocked down *dot-1.3* by RNAi specifically in the germline using the germline-specific *rde-1* rescuing strain (fig. S4K, S4L, table S4) and found that the germline-only inactivation of *dot-1.3* reduces the lifespan extension in the *lipl-4 Tg* worms (Fig. 4C and table S4). Moreover, rescuing *dot-1.3* expression in the germline restores the lifespan extension in the *lipl-4 Tg* worms with the *dot-1.3(lf)* mutant (Fig. 4D and table S2). Although no intestinal DOT-1.3 was detected in the CRISPR knock-in line, we still examined its potential involvement in regulating longevity. We found that *dot-1.3* RNAi knockdown specifically in the intestine does not suppress the lifespan extension in the *lipl-4 Tg* worms (fig. S4M, S4N and table S4). We further selectively restored *dot-1.3* expression in the intestine of the *lipl-4 Tg* worm with the *dot-1.3(lf)* mutant but observed no restoration of extended lifespan (fig. S4O and table S2). These results suggest that germline DOT-1.3 mediates the pro-longevity effect conferred by LIPL-4-induced lysosomal lipolysis.

Furthermore, we generated a transgenic strain to overexpress *dot-1.3* selectively in the germline (*dot-1.3 Tg*). We found that the *dot-1.3 Tg* worms exhibit an extended lifespan (Fig. 4E and table S2) and increase the H3K79me2 but not H3K79me3 level (Fig. 4F, fig. S4P and S4Q). These results suggest that, although *dot-1.3* knockdown is capable of suppressing the induction of both H3K79me2 and H3K79me3, its overexpression is only sufficient to induce H3K79me2, which may be related to the availability of methyl donors. Moreover, these results suggest that the induction of H3K79me2 alone is sufficient to promote longevity, and further support the importance of germline DOT-1.3 in regulating longevity.

### H3K79me2 transmission mediates transgenerational longevity

To examine whether H3K79me2 induction is transmitted across generations and mediates transgenerational longevity, we first conducted ChIP-seq analysis using anti-H3K79me2 antibodies and compared the T2 WT descendants from the *lipl-4 Tg* cross and the *dot-1.3 Tg* cross with WT controls. We found that the H3K79me2 occupancy in the genome is increased in both T2 WT descendants (Fig. 4G). Next, we used the *dot-1.3(lf)* mutant and confirmed that it suppresses the lifespan extension in the T2 WT descendants from the *lipl-4 Tg* cross (fig. S4R, S4S and table S1). We further knocked down *dot-1.3* by RNAi only during the larval stage of T1 WT descendants from the *lipl-4 Tg* worms and found this fully abrogated the lifespan extension in the T2 WT descendants (Fig. 4H and table S1). Moreover, the T2 WT descendants from the *dot-1.3 Tg* worms showed extended lifespan compared to WT controls (Fig. 4I). Together, these results demonstrate the importance of DOT-1.3-mediated H3K79 methylation in regulating transgenerational longevity.

We also found that germline HIS-71 requires K79 methylation for its pro-longevity effects. In the *dot-1.3(lf)* mutant, the germline-specific overexpression of *his-71* failed to extend lifespan (fig. S4T and table S2). Furthermore, we generated a transgenic strain to overexpress a mutant form of HIS-71 (K79A) that substitutes the lysine residue at 79 with an alanine which is unable to be methylated (*27*). We found that germline-specific overexpression of this K79A mutant form only led to a ∼9% change in lifespan (fig. S4U and table S2), which is much weaker than over 20% lifespan extension caused by the WT HIS-71 overexpression in the germline (fig. S3J and table S2). Additionally, we found that the lifespan extension in the *dot-1.3 Tg* worms is fully abrogated by the *H3.3* null mutant (fig. S4V and table S2), whereas the *his-71(lf)* mutant alone is insufficient (fig. S4W and table S2), suggesting that the germline DOT-1.3 may target other H3.3 variants in the absence of HIS-71. Nevertheless, the transgenerational pro-longevity effect induced by *dot-1.3 Tg* requires HIS-71 (Fig. 4J, table S1). Together, these findings uncover the previously unknown HIS-71-DOT-1.3 epigenetic axis in regulating transgenerational longevity.

### Lysosome-related metabolic signaling regulates transgenerational longevity

We further tested whether HIS-71 and DOT-1.3 also contribute to other longevity regulatory mechanisms, including reduction in insulin/IGF-1 signaling (IIS), mTOR signaling or mitochondrial electron transport chain (ETC) complexes, germline deficiency and caloric restriction (*28*). We have crossed the *his-71(lf)* and the *dot-1.3(lf)* mutant with the *daf-2(lf)* mutant (IIS reduction), the *raga-1(lf)* mutant (mTOR reduction), the *glp-1(lf)* mutant (germline deficiency) and the *eat-2(lf)* mutant (caloric restriction) or treated the *his-71(lf)* and the *dot-1.3(lf)* mutant with *cco-1* RNAi knockdown (ETC reduction). We found that the *dot-1.3(lf)* or the *his-71(lf)* mutant only suppresses the lifespan extension in the *raga-1(lf)* mutant but does not affect the pro-longevity effect conferred by the other mutants or the RNAi treatment (Fig. 5A, 5B, fig. S5A-H, tables S2 and S4). Consistently, we detected the transcriptional upregulation of *his-71* in the intestine of the *raga-1(lf)* mutant, which is reduced by the RNAi inactivation of the SKN-1 transcription factor (Fig. 5C). The H3K79me2 level is also elevated with the reduction of *raga-1* (Fig. 5D, fig. S5I and S5J).

**Fig. 5.**
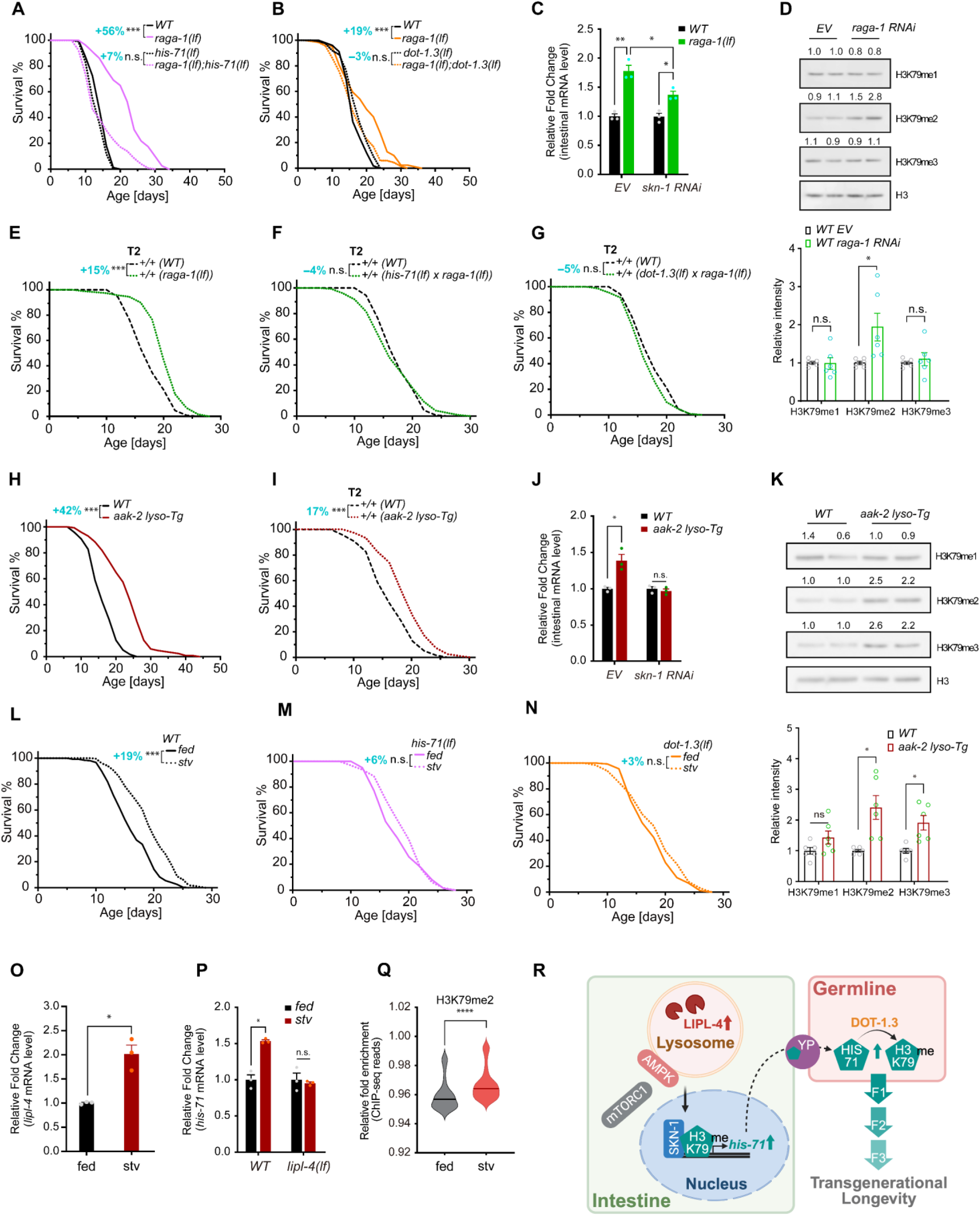
Lysosome-related metabolic signaling pathways regulate transgenerational longevity through HIS-71-DOT-1.3 axis. (**A**, **B**) Either the *his-71(lf)* mutant or the *dot-1.3(lf)* mutant reduces the lifespan extension caused by the *raga-1(lf)* mutant. (**C**) Intestinal upregulation of *his-71* transcription in the *raga-1(lf)* vs. WT worms is decreased by RNAi knockdown of *skn-1*. (**D**) RNAi inactivation of *raga-1* increases the H3K79me2 level. (**E-G**) T2 WT descendants from the *raga-1(lf)* mutant exhibit extended lifespan (E). This transgenerational lifespan extension is suppressed by the *his-71(lf)* mutant (F) and the *dot-1.3 (lf)* mutant (G). (**H**) Transgenic worms with intestinal lysosome-tethered AAK-2 overexpression (*aak-2 lyso-Tg*) show extended lifespan. (**I**) Transgenerational longevity is detected in T2 WT descendants from *aak-2 lyso-Tg*. (**J**) Intestinal upregulated *his-71* transcription is suppressed by RNAi knockdown of *skn-1* in *aak-2 lyso-Tg* vs. WT worms. (**K**) Western blots showing that H3K79me2 and me3 levels are elevated in *aak-2 lyso-Tg* vs. WT worms. (**L**-**N**) Starvation (stv)-induced lifespan extension (L) is abolished in the *his-71(lf)* (M) and the *dot-1.3(lf)* (N) mutants. (**O**, **P**) Starvation upregulates *lipl-4* and *his-71* transcription in WT worms but fails to upregulate *his-71* transcription in the *lipl-4(lf)* mutants. (**Q**) Starvation in WT worms elevates H3K79me2 deposition (±1 kb, TSS) when compared to fed condition. \*\*\*\**p* < 0.0001 (Wilcoxon test). Central lines denote median values of the data. (**R**) A model schematic illustrating transgenerational longevity driven by lysosome-related metabolic signaling pathways via the epigenetic axis HIS-71-DOT-1.3 across the intestine and germline. In Kaplan-Meier survival curves, n = 90/replicate; 3 biological replicates for (A, B, E-I, M, N) or 6 for (L); log-rank test followed by Fisher’s method: n.s., *p* > 0.05, \*\*\**p* < 0.001. Percentage of lifespan extension (lower vs. upper) is shown. Summary of lifespan replicates shown in tables S1, S2 and S4. In bar charts, error bars represent mean ±s.e.m.; 3 biological replicates for (C, J, O, P), or 6 for (D, K); n.s., *p* > 0.05, \**p* < 0.05 (unpaired t-test, Welch’s correction).

Rag GTPases, including RagA/B and RagC/D, facilitate the activation of mTOR complex 1 (mTORC1) by anchoring it onto the lysosome surface (*4*, *29*), and their inactivation causes the reduction of lysosomal mTORC1 signaling (*30*). In *C. elegans*, *raga-1* encodes RagA/B GTPase. We found that *raga-1* RNAi knockdown cannot further enhance the lifespan extension caused by *lipl-4 Tg* (fig. S5K and table S4), suggesting the interaction between these two lysosomal pro-longevity mechanisms. We then tested whether the reduction of lysosomal mTORC1 signaling could lead to transgenerational longevity. We crossed the *raga-1(lf)* mutant hermaphrodites with WT males, and those T2 WT descendants from the cross showed significant lifespan extension in comparison with WT controls (Fig. 5E and table S1). Furthermore, this lifespan extension was suppressed by the *his-71(lf)* and the *dot-1.3(lf)* mutant (Fig. 5F, 5G and table S1). These results support the role of lysosomal mTORC1 signaling in regulating transgenerational longevity through the HIS-71-DOT-1.3 epigenetic axis.

It was shown that AMPK docking at the lysosome competes with mTORC1, preventing its lysosomal activation (*3*). Recently, our lysosome-specific proteomic study revealed that AAK-2, a catalytic subunit of AMPK, is recruited to lysosomes in the *lipl-4 Tg* worms, contributing to their lifespan extension (*31*). We thus hypothesized that the lysosomal recruitment of AMPK in the intestine would result in a reduction in mTORC1 activity, thereby promoting transgenerational longevity as observed in WT descendants of the *lipl-4 Tg* and *raga-1 (lf)* worms. To test this, we generated a transgenic strain (*aak-2 lyso-Tg*) in which AAK-2 is tethered to lysosomes by fusion with LMP-1, a lysosome associated membrane protein, and then specifically overexpressed in the intestine. In parallel, a transgenic strain expressing a fluorescence protein mScarlet fused with LMP-1 in the intestine (*mScarlet lyso-Tg*) was generated as a control. The *aak-2 Tg* worms (*32*), in which *aak-2* is ubiquitously expressed without lysosomal tethering, were also included for analyses. We observed comparable lifespan extensions in the *aak-2 lyso-Tg* (42%) and *aak-2 Tg* (31%) worms, but no lifespan extension in the *mScarlet lyso-Tg* controls (Fig. 5H, fig. S5L, S5M, table S2).

We next examined the lifespans of their T2 WT descendants and found a 17% extension in T2 WT from the *aak-2 lyso-Tg* worms (Fig. 5I), but not in T2 WT from the *aak-2 Tg* worms (0%, fig. S5N and table S1). Similar to the findings in the *lipl-4 Tg* and *raga-1(lf)* worms, the *aak-2 lyso-Tg* worms exhibited upregulated intestinal *his-71* transcription in a SKN-1 dependent manner (Fig. 5J), as well as elevated H3K79 methylation (Fig. 5K, fig. S5O, S5P). Furthermore, RNAi inactivation of *dot-1.3* in the T1 WT descendants from *aak-2 lyso-Tg* abrogated their transgenerational pro-longevity effect (fig. S5Q and table S1). Together, these results suggest that lysosomal activation of AMPK promotes transgenerational longevity via the HIS-71-DOT-1.3 epigenetic axis.

Both lysosomal mTORC1 and AMPK signaling have been implicated in the metabolic response to starvation, a crucial environmental change that induces epigenetic reprogramming and lifespan extension (*4*, *6*, *33-35*). LIPL-4 is also transcriptionally upregulated upon fasting (*1*, *2*). To evaluate the contribution of the LIPL-4-mediated HIS-71-DOT-1.3 epigenetic axis to starvation-induced lifespan extension, we subjected the L1 larvae of worms to six days of starvation and assessed their adulthood lifespan following refeeding, with worms fed *Ad libitum* as the control. Consistent with previous findings (*35*), this starvation regimen extended the lifespan of WT worms by 19% (Fig. 5L and table S2); however, this effect was abolished in the *his-71(lf)* or the *dot-1.3(lf)* mutants (Fig. 5M, 5N and table S2) and was reduced to 10% in the *lipl-4(lf)* mutant (fig. S5R and table S2). Furthermore, we observed increased *lipl-4* and *his-71* transcriptional level and elevated H3K79me2 deposition across the genome in day-1 adult WT worms experiencing L1 larval starvation (Fig. 5O-Q). Notably, the transcriptional upregulation of *his-71* by starvation is fully suppressed in the *lipl-4(lf)* mutant (Fig. 5O). These results support the induction of LIPL-4-mediated HIS-71-DOT-1.3 epigenetic axis in response to starvation under physiological conditions.

## Discussion

Collectively, our data demonstrates that lysosomal metabolic pathways signal through the H3.3-H3K79me2 epigenetic axis to regulate transgenerational longevity (Fig. 5R). We propose that in response to environmental inputs like starvation, the induction of lysosomal acid lipase LIPL-4 leads to the lysosomal recruitment of AMPK and consequently reduced activation of mTORC1, which upregulates the transcription of the histone 3.3 variant HIS-71 through the transcription factor SKN-1. Intestine-induced HIS-71 is transported via the vitellogenin secretion and uptake pathway into the germline where it is post-translationally modified by the H3K79 methyltransferase DOT-1.3 and transmitted across generations to promote longevity. It is known that H3.3-enriched epigenomes exhibit distinct transcriptional responses, setting them apart from canonical histone H3-enriched epigenomes (*14*, *15*). The transition from H3.3 to H3-enriched epigenome restricts embryonic pluripotency and plasticity during *C. elegans* development (*21*). H3.3 variants can also assist in maintaining genome stability (*36-38*) and facilitate the repair of DNA damage (*39*, *40*). We thus anticipate that increased incorporation of H3.3 may help preserve a “youthful” epigenomic state during aging.

This study reveals a previously unknown cross-tissue epigenetic coordination in regulating transgenerational longevity. Sensitive to environmental changes, lysosomes serve as a vital hub to integrate metabolic signals and transcriptional responses cell-autonomously. Lysosomal metabolic signals, including AMPK and mTORC1, have been associated with longevity regulation across various species (*4*). Interestingly, lysosomes can transmit these signals cell non-autonomously from the soma to the germline by upregulation of histone H3.3 variants, which synergize with K79 methylation in the germline to carry longevity memory across multiple generations. Meanwhile, H3.3 variants are present in many cells, including the germline, and their local upregulation can also play a crucial role in promoting transgenerational longevity.

In this study, genetic evidence suggests that the yolk provision from the intestine to oocytes facilitates the transportation of HIS-71/H3.3. In *Drosophila*, maternal histone proteins are sequestered in lipid droplets and secreted from nurse cells to oocytes (*41*). Worm vitellogenins are homologous to human apolipoprotein B-100 and microsomal triglyceride transfer proteins, which are essential for the assembly, secretion, and circulation of very-low-density lipoproteins (*23*). In future research, it would be interesting to investigate the potential existence of histone trafficking and its biological significance in other organisms, as well as the transgenerational influences of AMPK and mTORC1 in mammals.

