## Supplementary figures for "Lysosomes Signal through Epigenome to Regulate Longevity across Generations"

#### fig. S1

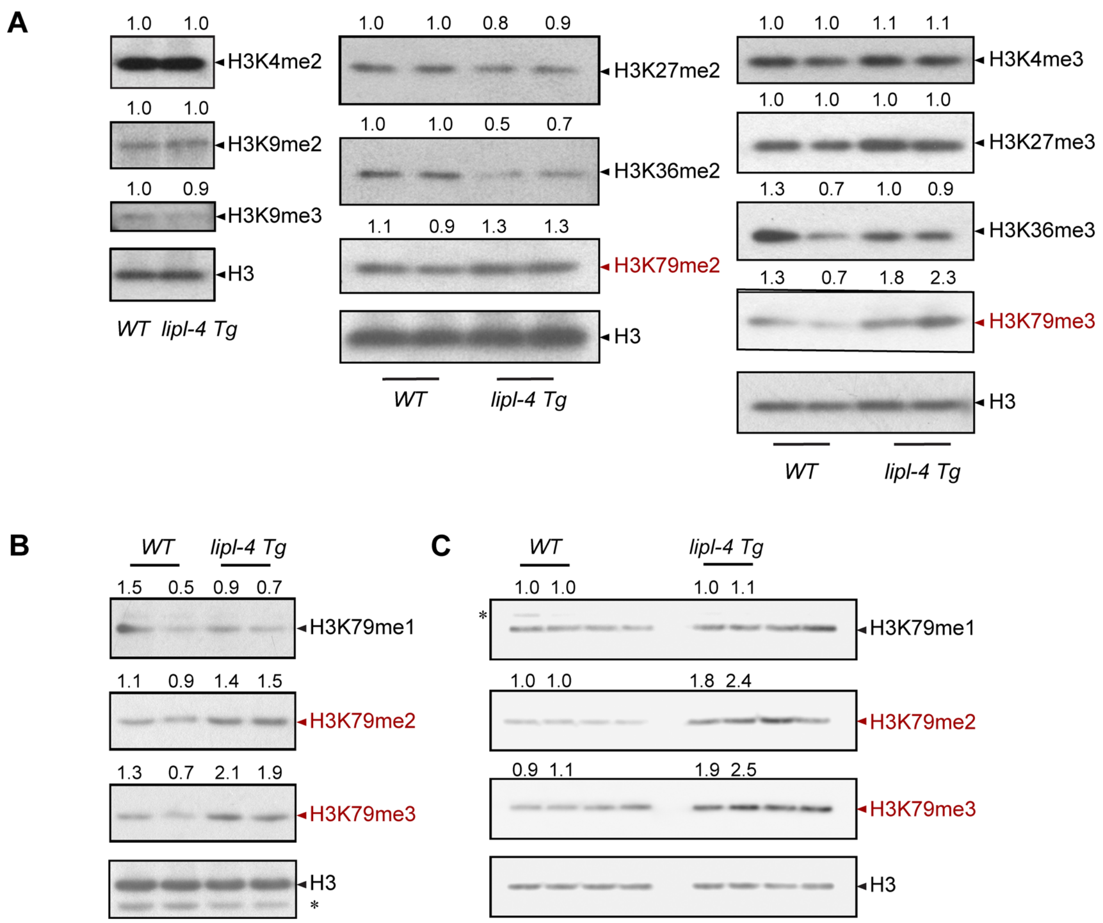


**fig. S1.** (**A-C**) Western blot images illustrating histone H3 PTM levels in *lipl-4 Tg* vs. WT worms. The asterisks indicate non-specific bands.

### fig. S2

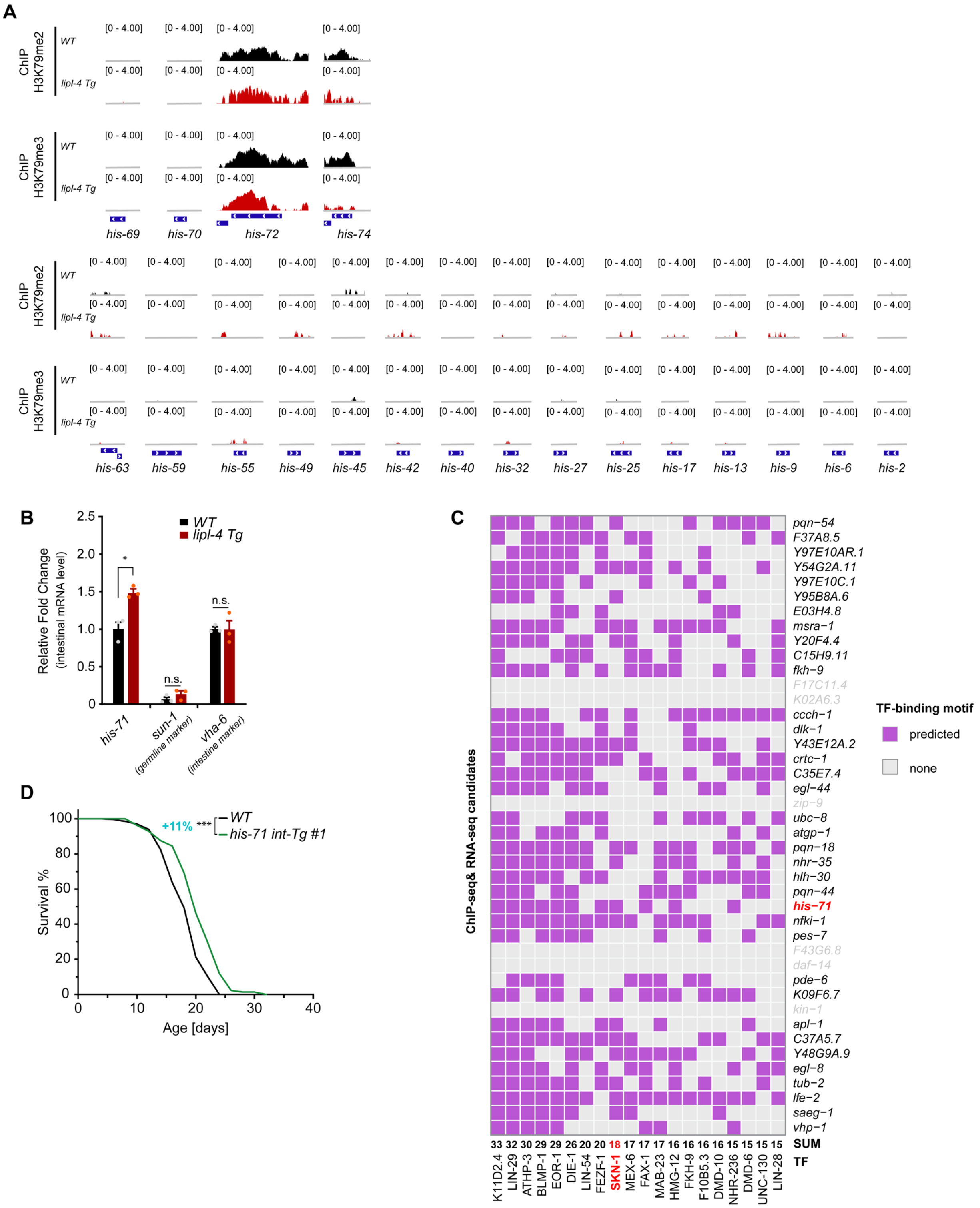


**fig. S2.** (**A**) Distribution of H3K79me2 and H3K79me3 marks at genes encoding H3.3 variants (*his-69*, *his-70*, *his-72*, *his-74*) and canonical H3 (*his-2, his-6, his-9, his-13, his-17, his-25, his-27, his-32, his-40, his-42, his-45, his-49, his-55, his-59, his-63*) in WT and *lipl-4 Tg* worms. The y-axis of ChIP-seq track is shown in a log scale (0-4.00). (**B**) Intestinal *his-71* transcription is upregulated in *lipl-4 Tg* vs. WT worms. Tissue-specific gene markers were included for validation: *sun-1* for the germline and *vha-6* for the intestine. Error bars represent mean ± s.e.m., n.s., $p > 0.05$, *$p < 0.05$ (unpaired t-test, Welch's correction). (**C**) Heatmap showing the predicted transcription factor (TF)-binding motifs (name at the bottom) within the promoter of candidate genes (name on the right) that are both enriched with H3K79me2 and H3K79me3 marks and transcriptionally upregulated (> 1.5-fold) in *lipl-4 Tg* vs. WT worms. Violet indicates the presence of a predicted TF-binding motif within the promoter, while gray signifies its absence. The "SUM" row presents the total number of candidate genes with the assigned TF-binding motif. SKN-1 and *his-71* are highlighted in red, and the genes with promoters shorter than 1 kb are colored in gray. The *Caenorhabditis elegans* TF motifs with experimentally verified preferred binding sequences were sourced from the CisBP database v2.00. The software tool FIMO was utilized to scan and identify motif positions within 1000 bp promoter regions, based on a statistical cutoff of $p < 1.0e\text{-}4$. (**D**) Intestine-specific overexpression of *his-71* (*his-71 int-Tg*) extends lifespan. With n = 90/replicate, 3 biological replicates; log-rank test followed by Fisher's method: ***$p < 0.001$. Percentage of lifespan extension (lower vs. upper) is labeled. Summary of lifespan replicates shown in table S2.

#### fig. S3

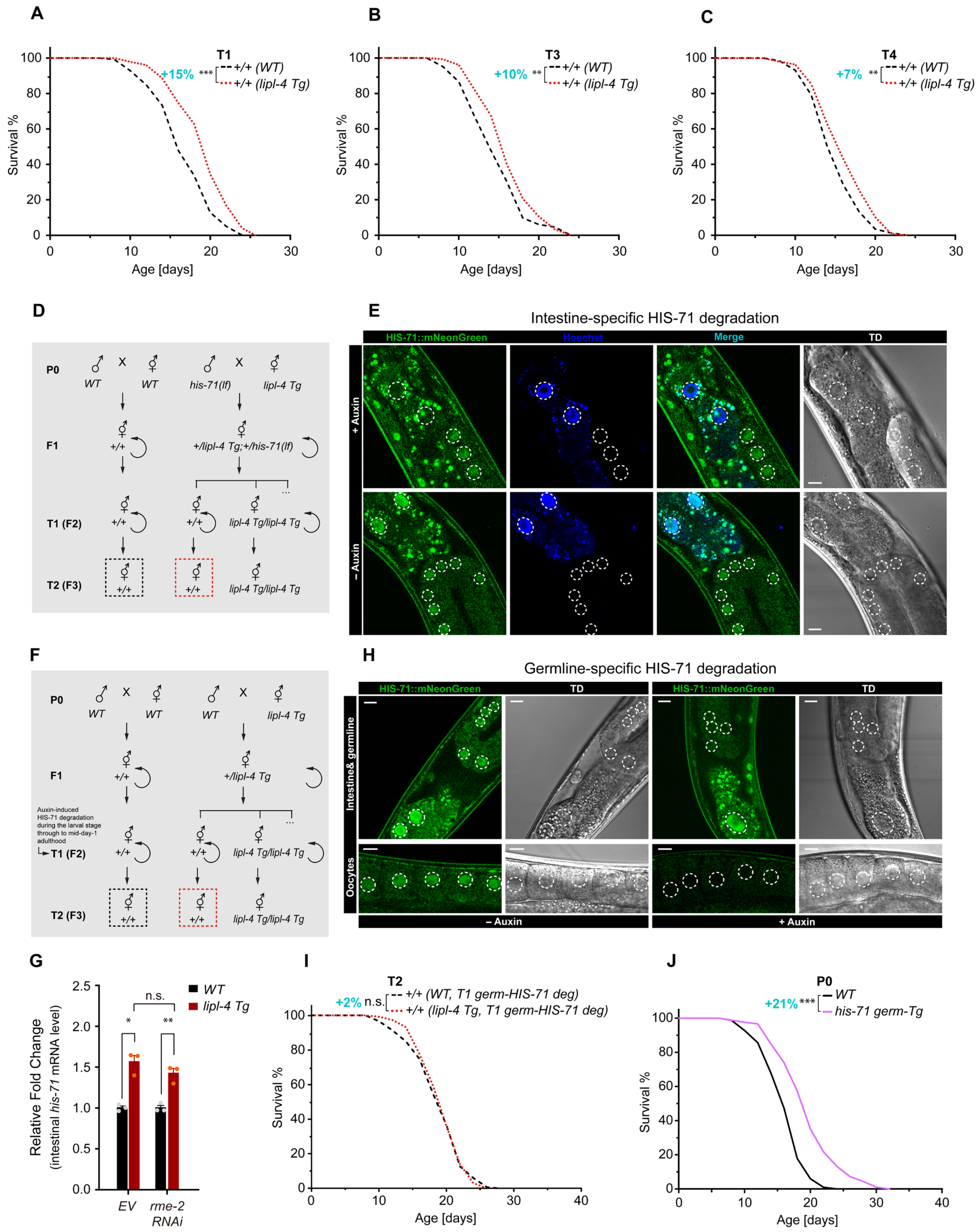
A
T1
+/+ (WT)
+/+ (lipl-4 Tg)
+15% ***
Survival %
Age [days]
B
T3
+/+ (WT)
+/+ (lipl-4 Tg)
+10% **
C
T4
+/+ (WT)
+/+ (lipl-4 Tg)
+7% **
D
P0
WT
his-71(lf)
lipl-4 Tg
F1
+/+
+/lipl-4 Tg;+/his-71(lf)
T1 (F2)
lipl-4 Tg/lipl-4 Tg
T2 (F3)
E
Intestine-specific HIS-71 degradation
HIS-71::mNeonGreen
Hoechst
Merge
TD
+ Auxin
– Auxin
F
+/lipl-4 Tg
Auxin-induced HIS-71 degradation during the larval stage through to mid-day-1 adulthood
H
Germline-specific HIS-71 degradation
Intestine& germline
Oocytes
G
Relative Fold Change (intestinal his-71 mRNA level)
WT
lipl-4 Tg
n.s.
EV
rme-2 RNAi
I
T2
+2% n.s.
+/+ (WT, T1 germ-HIS-71 deg)
+/+ (lipl-4 Tg, T1 germ-HIS-71 deg)
J
P0
+21% ***
WT
his-71 germ-Tg

#### fig. S3

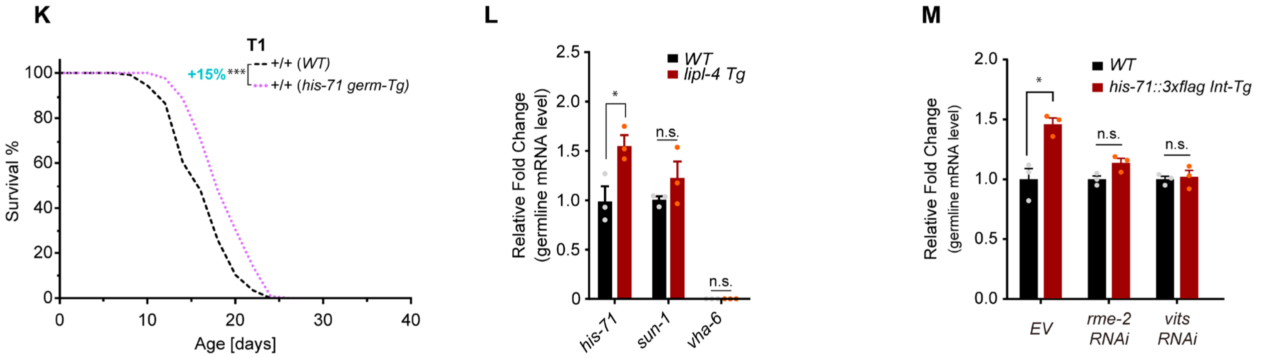


**fig. S3.** (**A**-**C**) Lifespans of WT descendants originating from *lipl-4 Tg* and WT worms across generations. (**D**) Experimental design for genetic crosses and tracking of subsequent generations. T2 WT worms (framed with dotted boxes) were used for lifespan analyses. (**E, H**) Confocal images showing auxin-induced degradation of HIS-71 specifically in the intestine (E) and germline (H). Green, endogenous HIS-71 tagged with mNeonGreen; blue, nuclei stained with Hoechst. Merged images show fluorescence overlay. Transmission Detection (TD) imaging provides morphological context. Auxin-treated (+auxin) or non-auxin treated (-auxin) are marked. Dotted lines encircle nuclei. Scale bar = 10 µm. In (H), top: intestine and germline; bottom: oocytes in focus. (**F**) Scheme for genetic crosses, generation tracking, and tissue-specific HIS-71 degradation. Lifespan analysis on T2 WT descendants (framed with dotted boxes). (**G**) RNAi knockdown of *rme-2* does not suppress the transcriptional upregulation of *his-71* in the intestine caused by *lipl-4 Tg*. (**I**) Germline degradation of HIS-71 in the T1 generation (T1 germ-HIS-71 deg) abolishes the transgenerational longevity in T2 WT from *lipl-4 Tg*. (**J**, **K**) Germline overexpression of *his-71* (*his-71 germ-Tg*) promotes longevity in the P0 generation and the T1 WT progeny. (**L**) Germline *his-71* transcription is upregulated in *lipl-4 Tg* vs. WT worms. (**M**) Intestinal overexpression of *his-71::3×flag* upregulates endogenous *his-71* transcription in the germline, which is suppressed by RNAi inactivation of *rme-2* or vitellogenin genes (*vits*). In (A-C, I-K), n = 90/replicate, 3 biological replicates; log-rank test followed by Fisher's method: n.s., $p > 0.05$, $^{**}p < 0.01$, $^{***}p < 0.001$. Percentage of lifespan extension (lower vs. upper) is labeled. Summary of lifespan replicates shown in tables S1 and S2. In (G, L, M), error bars represent mean ± s.e.m., n.s., $p > 0.05$, $^{*}p < 0.05$, $^{**}p < 0.01$ (unpaired t-test, Welch's correction).

fig. S4

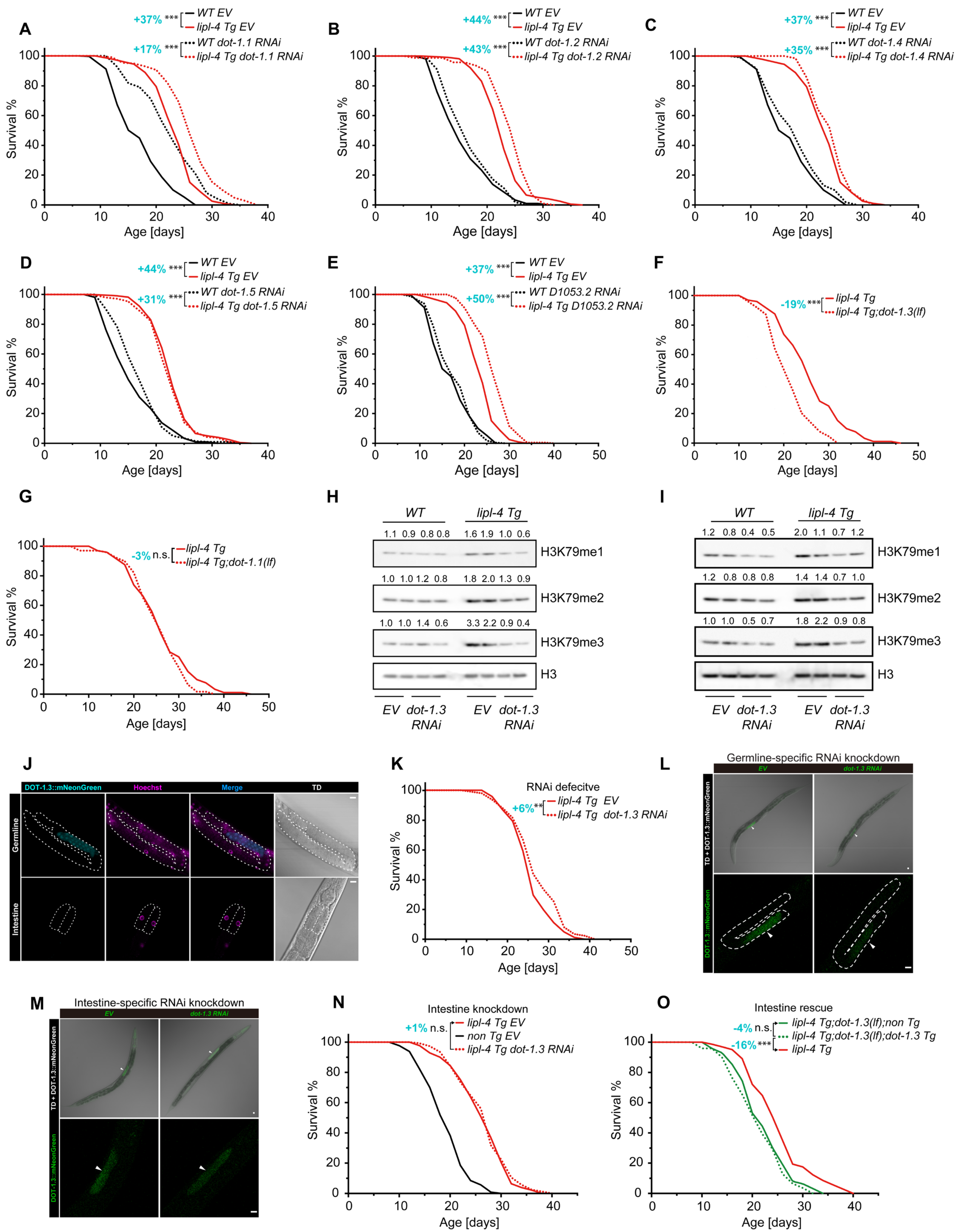
A
+37% *** WT EV
lipl-4 Tg EV
+17% *** WT dot-1.1 RNAi
lipl-4 Tg dot-1.1 RNAi
Survival %
Age [days]
B
+44% *** WT EV
lipl-4 Tg EV
+43% *** WT dot-1.2 RNAi
lipl-4 Tg dot-1.2 RNAi
C
+37% *** WT EV
lipl-4 Tg EV
+35% *** WT dot-1.4 RNAi
lipl-4 Tg dot-1.4 RNAi
D
+44% *** WT EV
lipl-4 Tg EV
+31% *** WT dot-1.5 RNAi
lipl-4 Tg dot-1.5 RNAi
E
+37% *** WT EV
lipl-4 Tg EV
+50% *** WT D1053.2 RNAi
lipl-4 Tg D1053.2 RNAi
F
-19% *** lipl-4 Tg
lipl-4 Tg;dot-1.3(lf)
G
-3% n.s. lipl-4 Tg
lipl-4 Tg;dot-1.1(lf)
H
WT
lipl-4 Tg
1.1 0.9 0.8 0.8 1.6 1.9 1.0 0.6
H3K79me1
1.0 1.0 1.2 0.8 1.8 2.0 1.3 0.9
H3K79me2
1.0 1.0 1.4 0.6 3.3 2.2 0.9 0.4
H3K79me3
H3
EV dot-1.3 RNAi EV dot-1.3 RNAi
I
WT
lipl-4 Tg
1.2 0.8 0.4 0.5 2.0 1.1 0.7 1.2
H3K79me1
1.2 0.8 0.8 0.8 1.4 1.4 0.7 1.0
H3K79me2
1.0 1.0 0.5 0.7 1.8 2.2 0.9 0.8
H3K79me3
H3
EV dot-1.3 RNAi EV dot-1.3 RNAi
J
DOT-1.3::mNeonGreen
Hoechst
Merge
TD
Germline
Intestine
K
RNAi defecitve
+6% ** lipl-4 Tg EV
lipl-4 Tg dot-1.3 RNAi
L
Germline-specific RNAi knockdown
EV
dot-1.3 RNAi
TD + DOT-1.3::mNeonGreen
DOT-1.3::mNeonGreen
M
Intestine-specific RNAi knockdown
EV
dot-1.3 RNAi
TD + DOT-1.3::mNeonGreen
DOT-1.3::mNeonGreen
N
Intestine knockdown
+1% n.s. lipl-4 Tg EV
non Tg EV
lipl-4 Tg dot-1.3 RNAi
O
Intestine rescue
-4% n.s. lipl-4 Tg;dot-1.3(lf);non Tg
lipl-4 Tg;dot-1.3(lf);dot-1.3 Tg
-16% *** lipl-4 Tg

**fig. S4**

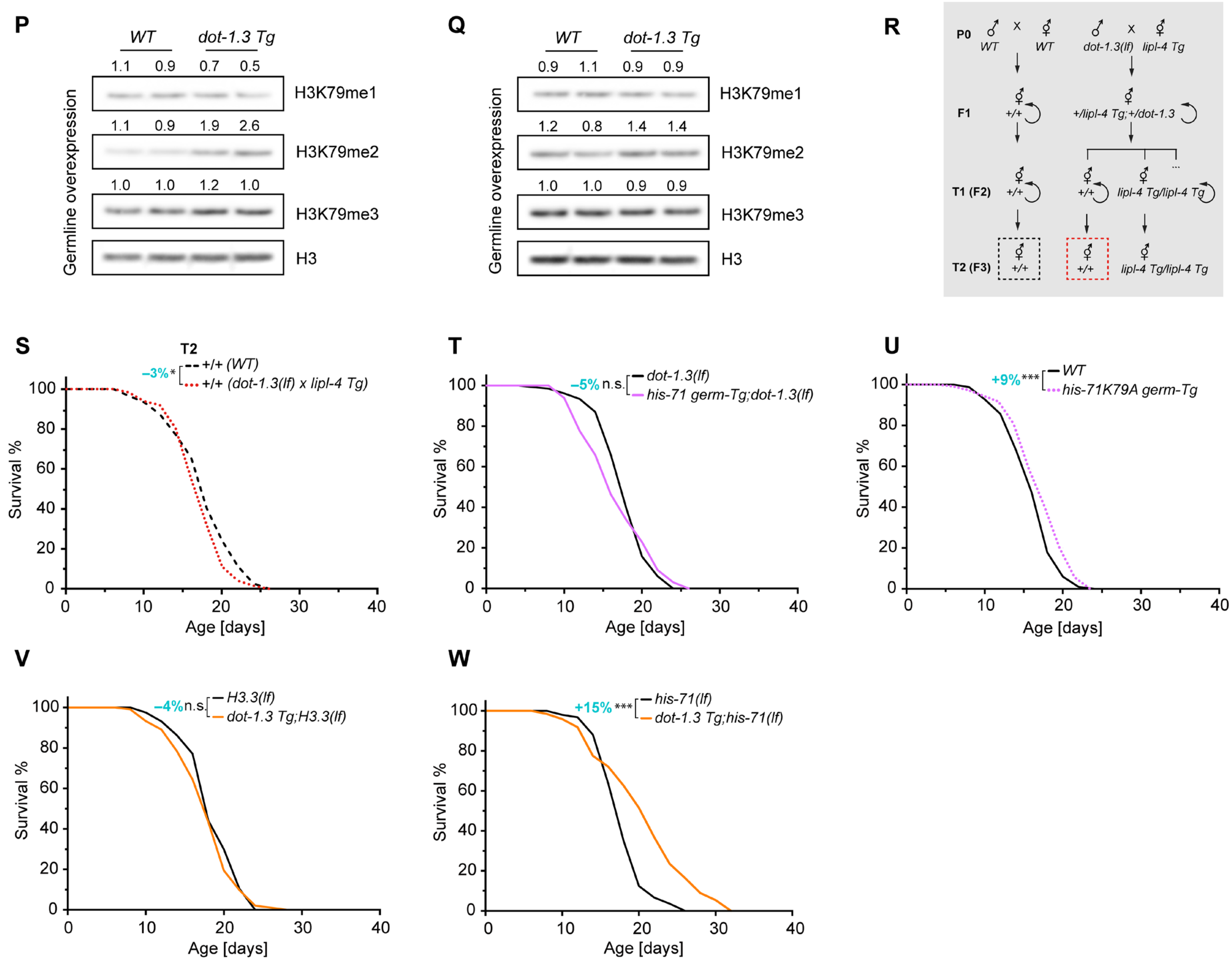


**fig. S4.** (**A-E**) The effects of RNAi inactivation of H3K79 methyltransferase encoding genes, *dot-1.1* (A), *dot-1.2* (B), *dot-1.4* (C), *dot-1.5* (D), *D1053.2* (E) on the lifespans of WT and *lipl-4 Tg* worms. (**F**, **G**) The *dot-1.3(lf)* but not the *dot-1.1(lf)* mutant decreases the lifespan of the *lipl-4 Tg* worms. (**H**, **I**) Western blot replicates on worms with *dot-1.3* RNAi knockdown. (**J**) Confocal images showing DOT-1.3::mNeonGreen fluorescent signals detected in the germline using a CRISPR knock-in line. Cyan, DOT-1.3::mNeonGreen; magenta, Hoechst stained nuclei. TD, Transmission Detection. Top: the dotted lines encircling the germline; bottom: the dotted lines encircling the intestine cells. Scale bar = 10 µm. (**K**) In the *rde-1* null mutant where RNAi is defective, *dot-1.3* RNAi fails to reduce the lifespan of *lipl-4 Tg* worms. (**L**) Validation of germline-specific knockdown of *dot-1.3* using worms carrying endogenous DOT-1.3::mNeonGreen. Top: merged images showing the overlap of mNeonGreen fluorescence and TD imaging (20× objective); bottom: enlarged fluorescent images with the germline encircled. Scale bar = 10 µm. (**M**) Intestine-specific knockdown of *dot-1.3* does not reduce the germline fluorescent signal from worms carrying DOT-1.3::mNeonGreen. Top: merged images showing the overlap of mNeonGreen fluorescence and TD imaging (20× objective); bottom: enlarged fluorescent images illustrating the mNeonGreen fluorescence in the germline. Scale bar = 10 µm. (**N**) Intestine-specific knockdown of *dot-1.3* does not shorten the lifespan of the *lipl-4 Tg* worms. (**O**) Restoration of *dot-1.3* expression in the intestine of *lipl-4 Tg;dot-1.3(lf)* mutant does not restore lifespan to the level of *lipl-4 Tg* alone. (**P**, **Q**) Western blot replicates on worms with germline-specific *dot-1.3* overexpression. (**R**) Scheme representing genetic crosses and transgenerational groups (framed with dotted boxes) used in lifespan analyses. (**S**) T2 WT descendants from the cross between *lipl-4 Tg* hermaphrodites and *dot-1.3(lf)* males do not show lifespan extension. (**T**) In the *dot-1.3(lf)* mutant background, germline-specific overexpression of *his-71* fails to extend lifespan. (**U**) Germline-specific overexpression of the HIS-71(K79A) mutant form results in negligible lifespan extension (< 10%). (**V**) In the H3.3(null) mutant background, *dot-1.3 Tg* fails to extend lifespan. (**W**) The *his-71(lf)* mutant alone does not suppress the lifespan extension in the *dot-1.3 Tg* worms. In Kaplan-Meier survival curves, n = 60-90 worms (A-E) or n = ~90 worms (F, G, K, N, O, S-W) per replicate, 3 biological replicates; log-rank test followed by Fisher's method: n.s., $p > 0.05$, *$p < 0.05$, **$p < 0.01$, ***$p < 0.001$. Percentage of lifespan extension (lower vs. upper, or indicated by arrows) is labeled. Summary of lifespan replicates shown in tables S1, S2 and S4.

fig. S5

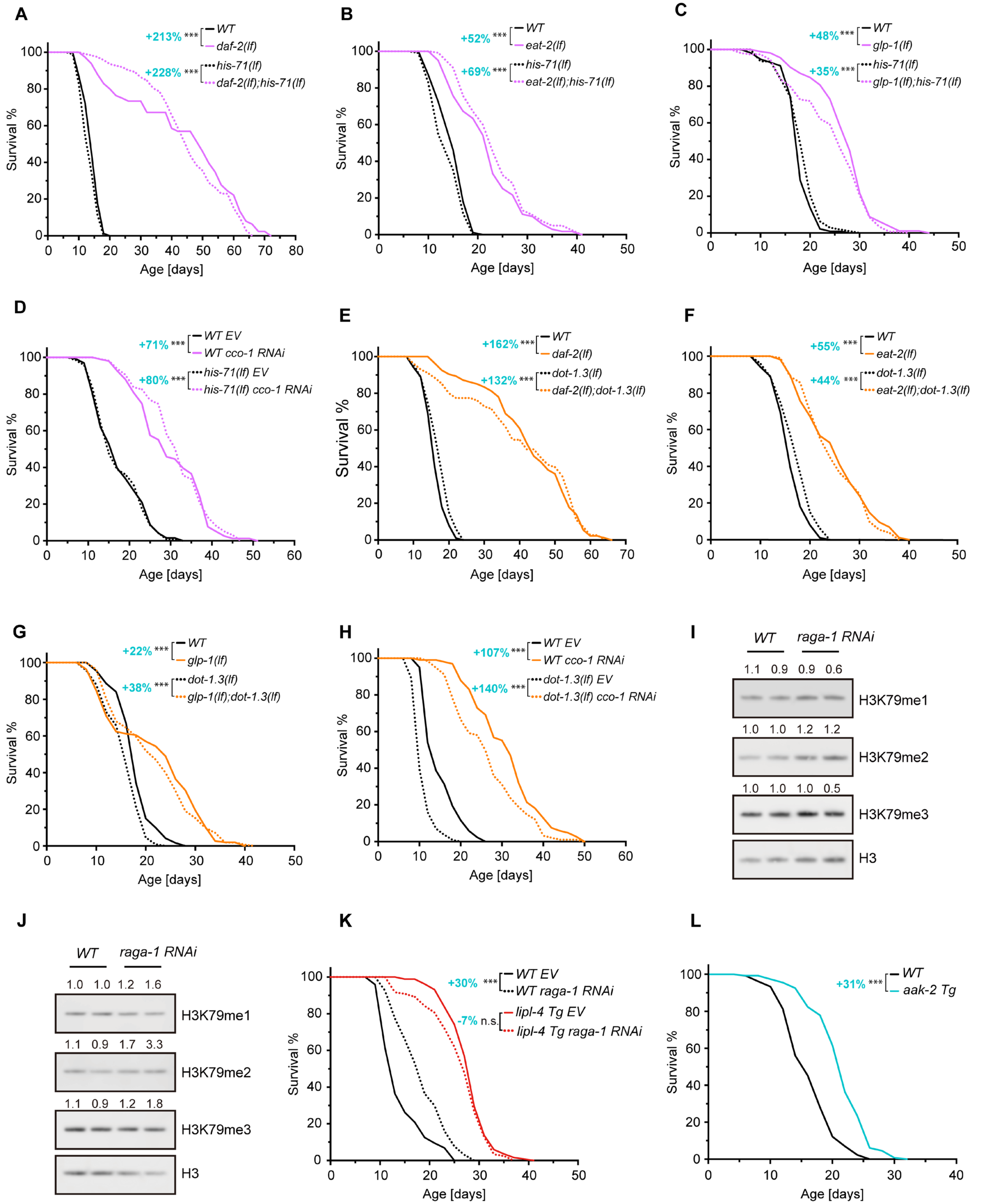
A
+213% *** WT
daf-2(lf)
+228% *** his-71(lf)
daf-2(lf);his-71(lf)
Survival %
Age [days]
B
+52% *** WT
eat-2(lf)
+69% *** his-71(lf)
eat-2(lf);his-71(lf)
Survival %
Age [days]
C
+48% *** WT
glp-1(lf)
+35% *** his-71(lf)
glp-1(lf);his-71(lf)
Survival %
Age [days]
D
+71% *** WT EV
WT cco-1 RNAi
+80% *** his-71(lf) EV
his-71(lf) cco-1 RNAi
Survival %
Age [days]
E
+162% *** WT
daf-2(lf)
+132% *** dot-1.3(lf)
daf-2(lf);dot-1.3(lf)
Survival %
Age [days]
F
+55% *** WT
eat-2(lf)
+44% *** dot-1.3(lf)
eat-2(lf);dot-1.3(lf)
Survival %
Age [days]
G
+22% *** WT
glp-1(lf)
+38% *** dot-1.3(lf)
glp-1(lf);dot-1.3(lf)
Survival %
Age [days]
H
+107% *** WT EV
WT cco-1 RNAi
+140% *** dot-1.3(lf) EV
dot-1.3(lf) cco-1 RNAi
Survival %
Age [days]
I
WT raga-1 RNAi
1.1 0.9 0.9 0.6
H3K79me1
1.0 1.0 1.2 1.2
H3K79me2
1.0 1.0 1.0 0.5
H3K79me3
H3
J
WT raga-1 RNAi
1.0 1.0 1.2 1.6
H3K79me1
1.1 0.9 1.7 3.3
H3K79me2
1.1 0.9 1.2 1.8
H3K79me3
H3
K
+30% *** WT EV
WT raga-1 RNAi
-7% n.s. lipl-4 Tg EV
lipl-4 Tg raga-1 RNAi
Survival %
Age [days]
L
+31% *** WT
aak-2 Tg
Survival %
Age [days]

### fig. S5

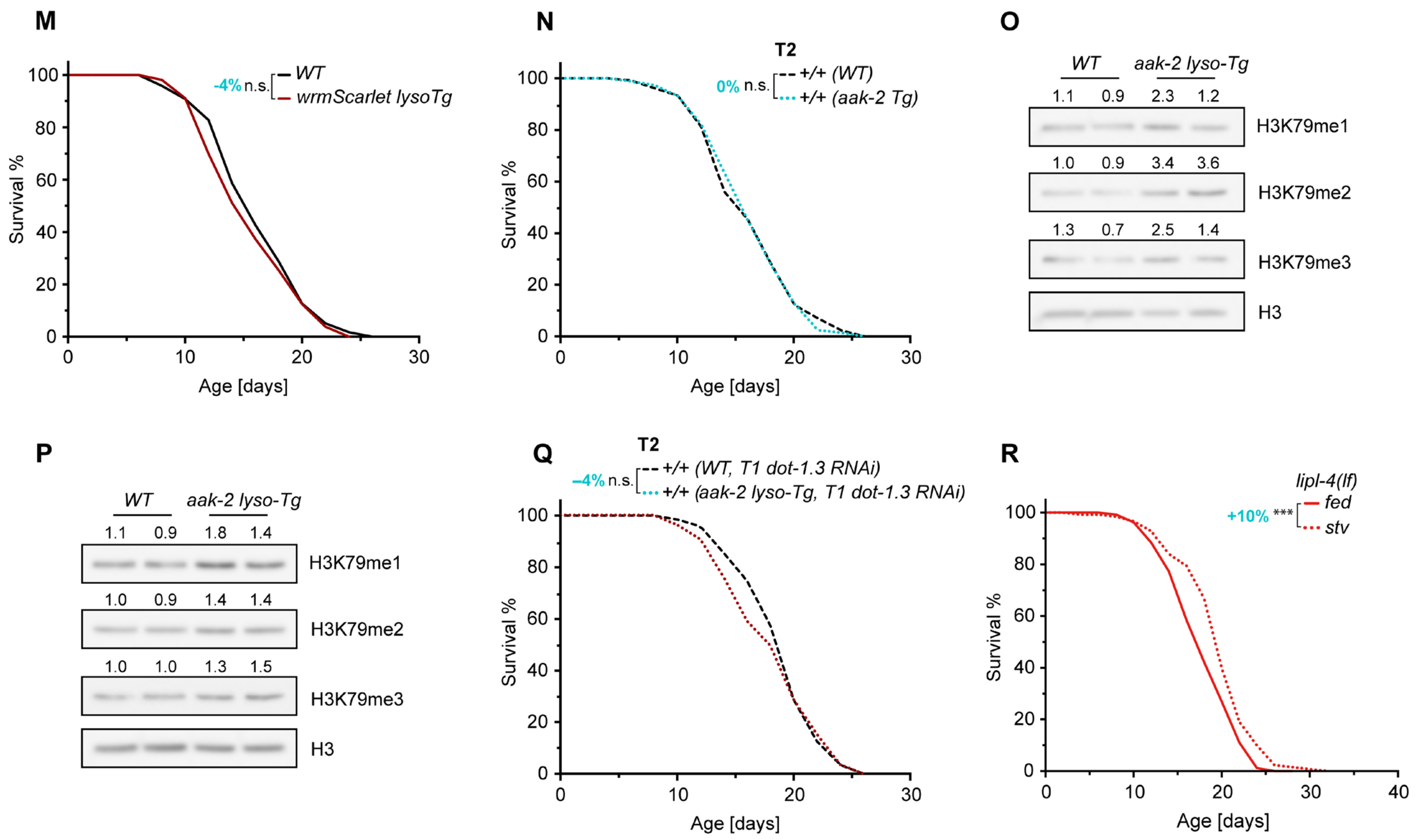


**fig. S5.** (**A**-**H**) The *his-71(lf)* (A-D) or the *dot-1.3(lf)* (E-H) mutant does not diminish the pro-longevity effects associated with the *daf-2(lf)*, *glp-1(lf)*, or *eat-2(lf)* mutant, or with the *cco-1* RNAi knockdown. (**I**, **J**) Western blot replicates on worms with *raga-1* RNAi knockdown. (**K**) RNAi inactivation of *raga-1* does not further enhance the lifespan extension caused by *lipl-4 Tg*. (**L**) Overexpression of *aak-2* without lysosomal tethering extends lifespan. (**M**) Transgenic worms overexpressing lysosome-tethered *wrmScarlet* in the intestine (*wrmScarlet lyso-Tg*) exhibit a lifespan comparable to WT worms. (**N**) Overexpression of *aak-2* without lysosomal tethering does not promote transgenerational longevity. (**O**, **P**) Western blot replicates on *aak-2 lyso-Tg* worms. (**Q**) Knockdown of *dot-1.3* in T1 WT descents of *aak-2 lyso-Tg* worms abolishes the transgenerational pro-longevity effect in T2 WT progeny. (**R**) Lifespan extension induced by starvation is reduced from 19% (WT) to 10% by the *lipl-4(lf)* mutant. In Kaplan-Meier survival curves, n = ~90/replicate, 3 biological replicates; log-rank test followed by Fisher's method: n.s., $p > 0.05$, ***$p < 0.001$. Percentage of lifespan extension (lower vs. upper as indicated) is labeled. Summary of lifespan replicates shown in tables S1, S2 and S4.
